# CMU Array: A 3D Nano-Printed, Customizable Ultra-High-Density Microelectrode Array Platform

**DOI:** 10.1101/742346

**Authors:** Mohammad Sadeq Saleh, Sandra M. Ritchie, Mark A. Nicholas, Rriddhiman Bezbaruah, Jay W. Reddy, Maysamreza Chamanzar, Eric A. Yttri, Rahul P. Panat

## Abstract

Microelectrode arrays (MEAs) provide the means to record electrophysiological activity fundamental to both basic and clinical neuroscience (e.g. brain-computer interfaces). Despite recent advances, current MEAs have significant limitations – including low recording density, fragility, expense, and the inability to optimize the probe to individualized study or patient needs. Here we address the technological limitations through the utilization of the newest developments in 3D nanoparticle printing.^1^ Our ‘CMU Arrays’ possess previously impossible electrode densities (> 6000 channels/cm^2^) with tip diameters as small as 10 μm. Most importantly, the probes are entirely customizable owing to the adaptive manufacturing process. Any combination of individual shank lengths, impedances, and layouts are possible. This is achieved in part via our new multi-layer, multi material, custom 3D-printed circuit boards, a fabrication advancement in itself. This device design enables new experimental avenues of targeted, large-scale recording of electrical signals from a variety of biological tissues.

## Introduction

Three-dimensional microelectrode arrays (MEAs) consisting of insulated, electrically conducting shanks are critical for a wide range of biological applications.^2-3^ They form the cornerstone of neuroscience and serve as the basis for the transformational field of brain-computer interfaces. An MEA’s success is dependent upon its sampling ability, a function of both the electrode density and ability to optimally target the brain regions of interest. While the current fabrication methods have achieved significant advances in recording density^4-5^ due to the developments in the micro electro-mechanical systems (MEMS) fabrication techniques,^6-10^ they are limited in their volumetric electrode densities, difficult to customize, and are implemented with notoriously brittle silicon (Si) materials. Because the “Utah arrays” are fabricated from metallized silicon using techniques such as dicing, etching, and photo or electron beam lithography, the pitch (distance between adjacent shanks) for such an array construction is limited by the kerf width of the dicing saw, limiting the recording density to a few hundred shanks per square centimeter. ^6, 11^ Further, the shank height variation is limited to uniformly varying lengths. In another approach, multiple, high-density in-plane electrodes can be fabricated on a single silicon or Printed Circuit Board (PCB) shank.^12^ By stacking multiple such shanks, a two or three-dimensional device can be built with 400-700 μm pitch, but the distance between the stacked shanks is limited by assembly and bonding space requirements.^13-14^ Recently, CMOS technology has enabled silicon-based Neuropixel probes, but the targeted areas are limited to what can be reached by a single linear shank, and fewer than 400 of the possible channels along this shank can be used at any time.^15^

The next generation of tools for electrophysiological recording must overcome the limiting factors above.^16^ Additionally, advances in the rapidly expanding field of neuroscience would greatly benefit from tailored probes specific to the study or patient. We note that the fabrication methods for MEAs have followed the trends in the semiconductor industry – moving from microwires to lithography. Recently, 3D nanoparticle printing has emerged as a new method to fabricate electronic devices that either complements lithography or fills in the critical length-scale gaps left by current methods.^17^ Microelectronics fabrication by inkjet^18^ and aerosol jet^19^ nanoparticle printing offers the freedom to use the target material in nano-dispersion form. This process enables rapid changes to the device layouts and the use of a variety of substrates to fabricate devices on, including flexible polymers. Flexible substrates can be used to create electrodes for uneven or moving surfaces, ideal for nerve-on-a-chip platforms^20^ or interfacing with cardiac tissue. The aerosol jet printing method can even print highly complex three-dimensional metallic lattices and spirals without any support materials.^1^

To this end, we developed a 3D nanoparticle printing system to open up an entirely new design space for three-dimensional bioelectronic devices. The objective was to exploit the rapid customization offered by 3D printing to demonstrate a new class of MEAs with high density and having arbitrary variation in shank heights, diameters, and routing. We demonstrate that the multi-scale, bottom-up printing method is able to assemble metal particles into three dimensional shanks with diameters of tens of micrometers and a length of several millimeters. Moreover, the printing method yields a significant cost and production time reduction. Further, we show that the high prototypability and reduction in the shank diameter leads to shank densities in excess of 6000 sites per cm^2^; an order of magnitude improvement over current methods.^11^ With these arrays, physiologists can optimize the MEA to target neural architectures ranging from dense populations in the cerebellum to multiple, layer-specific cortical ensembles - or both, simultaneously.

### Microelectrode Array Construction via 3D Printing

The construction of the customizable, acute ‘CMU Array’ microelectrodes was carried out using aerosol jet conformal printing method, which is a 3D nanoparticle printing technique. In this method, a metal nanoparticle dispersion is atomized using ultrasonic energy or pressurized gas to create a mist or aerosol consisting of microdroplets. Each microdroplet carries metal nanoparticles from the dispersion. The aerosol is driven to a nozzle using an inert gas, while a sheath gas focuses the aerosol droplets onto the substrate at a resolution of about 10 μm.^1^ Once the printing is complete, the ‘green’ shanks are heated in an oven to remove the binders and allow the sintering of the nanoparticles to form the conductive shanks.

Figures 1 shows the 3D nanoparticle printing method and the construction of the shanks for the three-dimensional MEAs. As shown in Fig. 1a, the process of forming the array involves stacking of the metal nanoparticle-containing droplets on top of each other to form the high aspect ratio shanks. The substrate is heated to 90-110 °C during printing, which allows solvent evaporation and rapid solidification of the droplets once they approach the substrate. The solidified droplets then form a base for the subsequent droplets, leading to the formation of a shank. The process is repeated at each shank location (See Methods section for details of the 3D printing process and Supplementary Video 1 for a video of the printing process). The droplet dispense is controlled by a shutter that can break and re-start the flow within 4 ms. Figure 1b demonstrates a series of printed shanks on alumina substrate, each with a tip diameter of 10 μm, a base diameter of 30 μm, and a pitch of 125 μm (density of 6400 shanks/cm^2^). The average aspect ratio along the length of the shanks in Fig. 1b is 50:1. Note that a 64-channel probe with 1mm long shanks takes under 90 minutes to construct by the 3D nanoparticle printing method.

**Figure 1.**
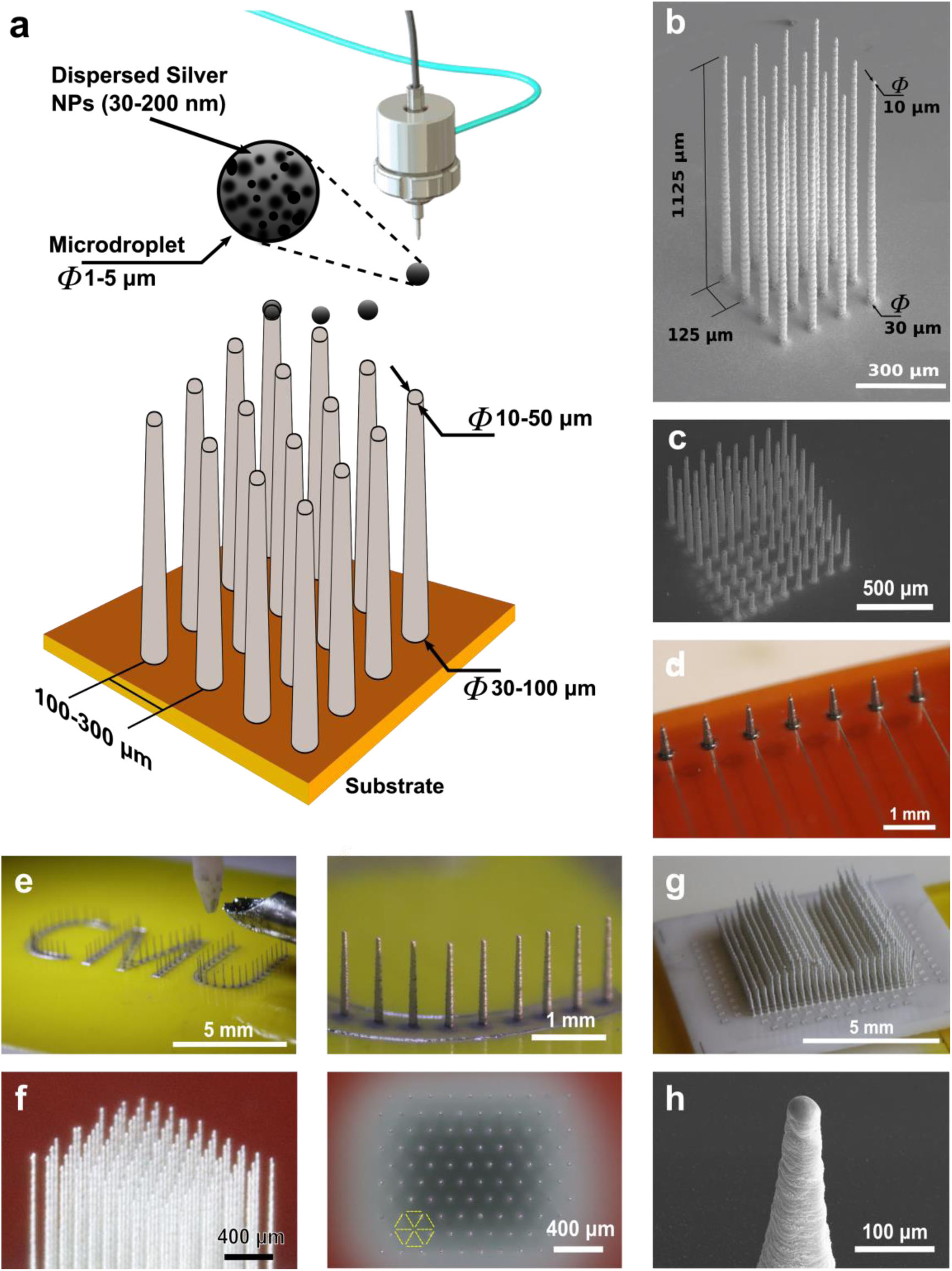
Microelectrode array Fabrication using 3D nanoparticle printing. (a) Schematic of the printing process, where aerosol droplets containing metal nanoparticles are stacked while instantaneously removing the solvent using the heat from the substrate. The long, narrow shanks are then sintered to remove the binders and create conductive paths to capture bioelectric signals. (b) Printed microelectrode array with 1.125 mm long shanks having an aspect ratio of 1:50 at a 125 μm grid. The substrate is alumina ceramic. (c) Printed shanks of different lengths and diameters (length=100-650 μm, diameters = 35-50 μm). Any combination of lengths and diameters is possible in the finalized probe. (d) Example of microelectrodes directly printed on a flexible polyimide (Kapton^®^) substrate. Wiring is also printed on the same substrate to connect the shanks to an external circuit. (e) Printed microelectrodes in an arbitrary pattern to resemble the Carnegie Mellon University abbreviation ‘CMU’, showing the extent of flexibility in design and fabrication by the 3D printing method in (a). A close-up of the shanks is shown on the right. (f) High density-high aspect ratio array printed in hexagonally packed pattern with 200 μm pitch distance. A 15% improvement in areal electrode density can be achieved by pattering the electrodes in hexagonal pattern rather than square packing. (g) A 512-electrode shank array with variable shank heights demonstrating the customization possible by this method. A time-lapse video of the construction of the probe is shown in Supplementary Video 2. (h) Examples demonstrating the control of the shank tips to facilitate their insertion and tolerance in biological tissue.

Because of the ease of customization afforded by computer aided design, this method of construction allows rapid changes to individual shank height and probe layout. Figure 1c shows silver shanks of varying heights within the same probe made by 3D nanoparticle printing. This feature will allow the capture of electrical signals from different depths within the brain or other biological tissue. Since printing can be done on any substrate, a wide range of rigid and flexible substrates can be used for the construction of the probe. Arrays may be printed on a flexible Kapton^®^ (polyimide) polymer substrate (Fig. 1d), enabling high-density, custom probes designed for use on curved or moving tissue. As a demonstration of the extent of flexibility in the design and fabrication of the shanks, we printed shanks bearing the CMU Array’s name (Fig. 1e). The shanks in this array are 1 mm tall with shank diameter ranging from 90 μm at the bottom to 30 μm at the top. To increase packing density by 15% over the traditional square array, CMU Array shanks can be arranged in a hexagonal pattern (Fig. 1f). Note that it is difficult if not impossible to create a hexagonal array pattern in traditional Utah arrays due to the limitations imposed by its manufacturing process.

Figure 1g demonstrates a 512-shank array with variable shank heights of 1.0, 1.5, and 2 mm, which demonstrates the possible customization of the probe. See Supplementary Video 2 for a time-lapse video of the construction of this probe. The AJ printing process also enables exquisite control of shaping the tip profile (Fig. 1h). For each of the shanks shown in Fig. 1, their angle to the vertical is within +/-1°. The results shown in Fig. 1 demonstrate that the 3D nanoparticle printing can be used to fabricate highly-customizable electrode arrays with high spatial densities. In the next section, we demonstrate probe structural properties, followed by custom electrical wiring, functionalization, and recording.

### Characterization of Nano-enabled Probe Properties

Electrode shanks must have the strength to penetrate the biological tissue of interest, such as the brain tissue or cardiac muscle. In addition, shanks should ideally strike a balance with ductility to tolerate outside forces without fracturing, including initial resistance from dura matter or user error. In the case of silicon probes, while Si is exceedingly strong, the extreme brittleness of Si can make the probes prone to breakage. Previously, we have demonstrated highly ductile metal pillars of a few tens of micrometers in diameter under a compressive load without significantly losing strength.^21^ We tested the high aspect ratio electrode shanks fabricated in this study (in an array of 3×3 shanks) and measured the load as a function of the displacement (Fig. 2a). We observed that for a displacement of the platen of up to 5%, the shanks show linear (elastic) deformation without breakage. The force required to bend the printed shanks shown was of the order of 0.3 N/shank (Supplementary Fig. 1). Note that in studies of probe robustness, the force required to penetrate agarose brain phantoms for a large, 200 μm diameter needle is of the order of 0.025 N.^22^ These shanks were further compressed without breakage but they enter a nonlinear (plastic) regime, indicating permanent deformation.

**Figure 2.**
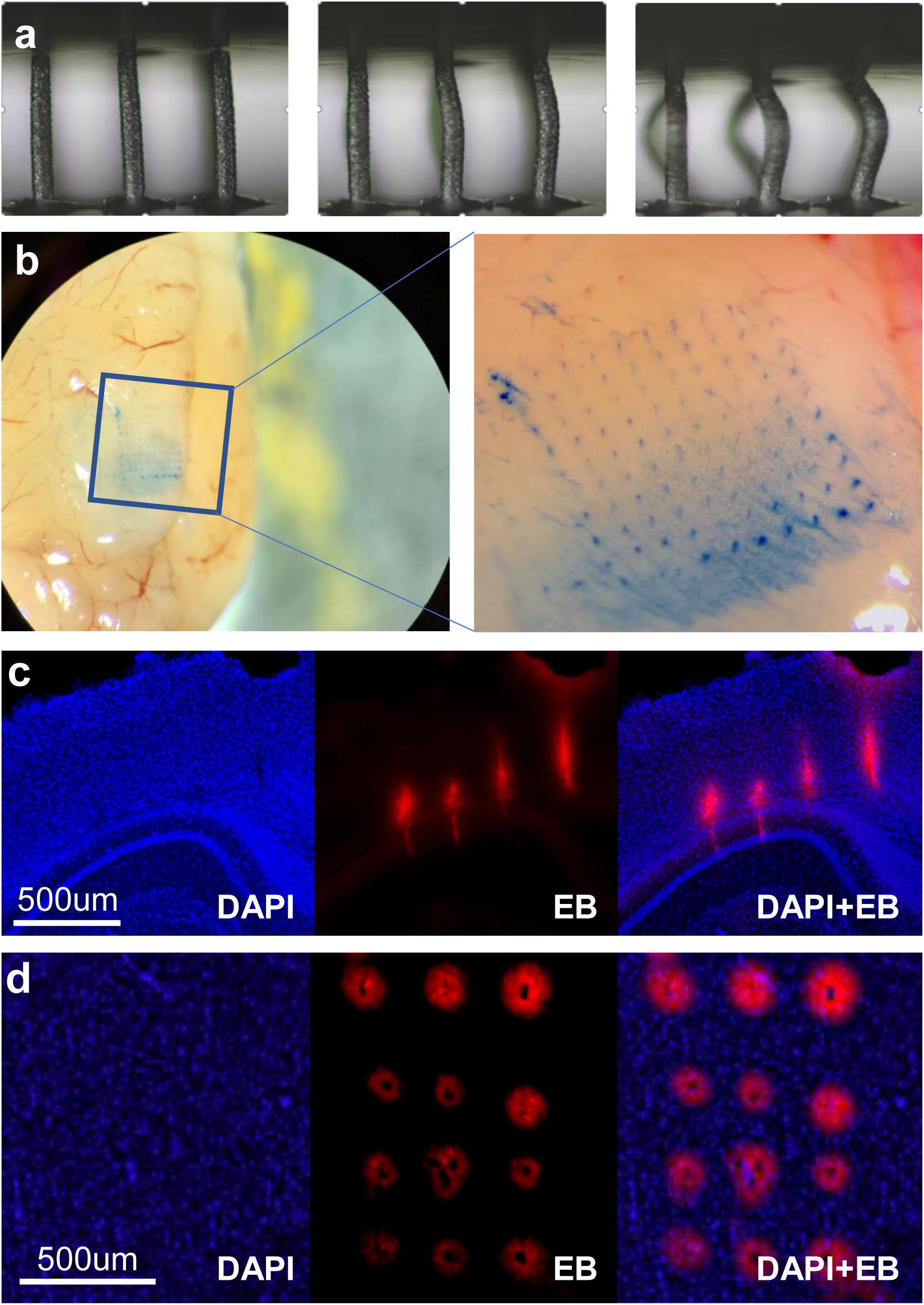
Mechanical properties of the probes and their insertion into mouse brain. (a) A 3×3 shank array under various forces of compression under a rigid platen. For a displacement of the rigid platen of up to 5%, the shanks show linear (elastic) deformation without breakage. The force-strain curve of this experiment is shown in Supplementary Fig. 1 and gives a maximum bending force of 0.3 N/shank. The shanks can be further compressed without breakage but they are in a nonlinear (plastic) space, indicating permanent deformation. (b) Evans blue dye left behind from the successful insertion of a dense (6400 shanks/cm^2^), 10×10 array of 20 μm diameter array with zoomed image (right). (c) Successful insertion test through dura in an anesthetized mouse without breaking the shanks nor gross damage to the brain (coronal slice though area V2, hippocampus; purple = DAPI nuclear stain, red = Evan’s blue). (d) Horizontal brain slice from insertion test into monkey cortex. Note the lack of damage or tearing caused by probe insertion in c,d.

Having achieved more than sufficient structural qualities, we next tested probe performance in tissue. We inserted a dense (6400 shanks/cm^2^), 10×10 array of 20 μm diameter, uniform height shanks into the cortex of an anesthetized mouse. Higher spatial density of probes, even with densities half that shown here, can lead to a ‘bed of nails’ effect in soft tissue, where the tissue is depressed rather than penetrated during insertion.^23^ In an attempt to overcome this undesirable effect, expensive accessory equipment are used, including vibration drives or a pneumatic insertion hammers. Arrays with staggered shank lengths are also often used to reduce the number of shanks penetrating at any given time. Owing to the small cross-sectional area and narrow tips, the 10×10 array of uniform shank length was capable of penetrating mouse brain with a basic, benchtop manual manipulator (Fig. 2b) at a rate of 0.2 mm/min. Histological examination of penetrations made with quite longer shank (length >2 mm, 15 μm diameter at tip, 75 μm at base, 25 shanks) also penetrate brain tissue without issue (Fig. 2c). Insertion test of this probe showed penetration through both mouse dura and brain, through area V2 to the hippocampus (Fig. 2c, right). Figure 2d shows the insertion of a probe into a macaque brain. In neither case did we find gross tissue damage, tearing, or other damage as acute probe slid through the brain tissue.

The highly-customizable 3D printed CMU Array platform has sufficient strength and ductility to penetrate biological tissue such as brain. Further, the ability to print sharp shank tips (Fig. 1h) further facilitates the entry of the shanks without inflicting tissue damage. At present, we are able to repeatedly penetrate brain with arrays possessing an extremely high shank density of 6400 sites/cm^2^ due to the method of construction and small cross-sectional area.

### Probe Functionalization and Recording of Action Potentials

In order to functionalize the probe, shanks were coated with a conformal insulating layer followed by exposure of the metal through selective removal of the polymer from the tips (Fig. 3). The shanks were first coated with a 3 μm insulating layer of biocompatable Parylene C polymer in a vacuum chamber using a chemical vapor deposition (CVD) process (Fig. 3b). The insulation was then removed selectively at the tip of the electrodes using a focused ion beam (FIB) cut with an exposed area in the range of 100-150 μm^2^ (Fig. 3c). The interface impedance of the electrodes can be improved by coating the tips of the shanks with electrically conducting polymers such as poly(3,4-ethylenedioxythiophene), also known as PEDOT (Fig. 3d). The electrochemical impedance of the electrode sites at the tip of each shank was then measured. We observed impedances in the range of 200-400 kΩ at a frequency of 1 kHz (see Supplementary Fig. 2). These impedance values are within the desired range for extracellular recording.^13, 24^

**Figure 3.**
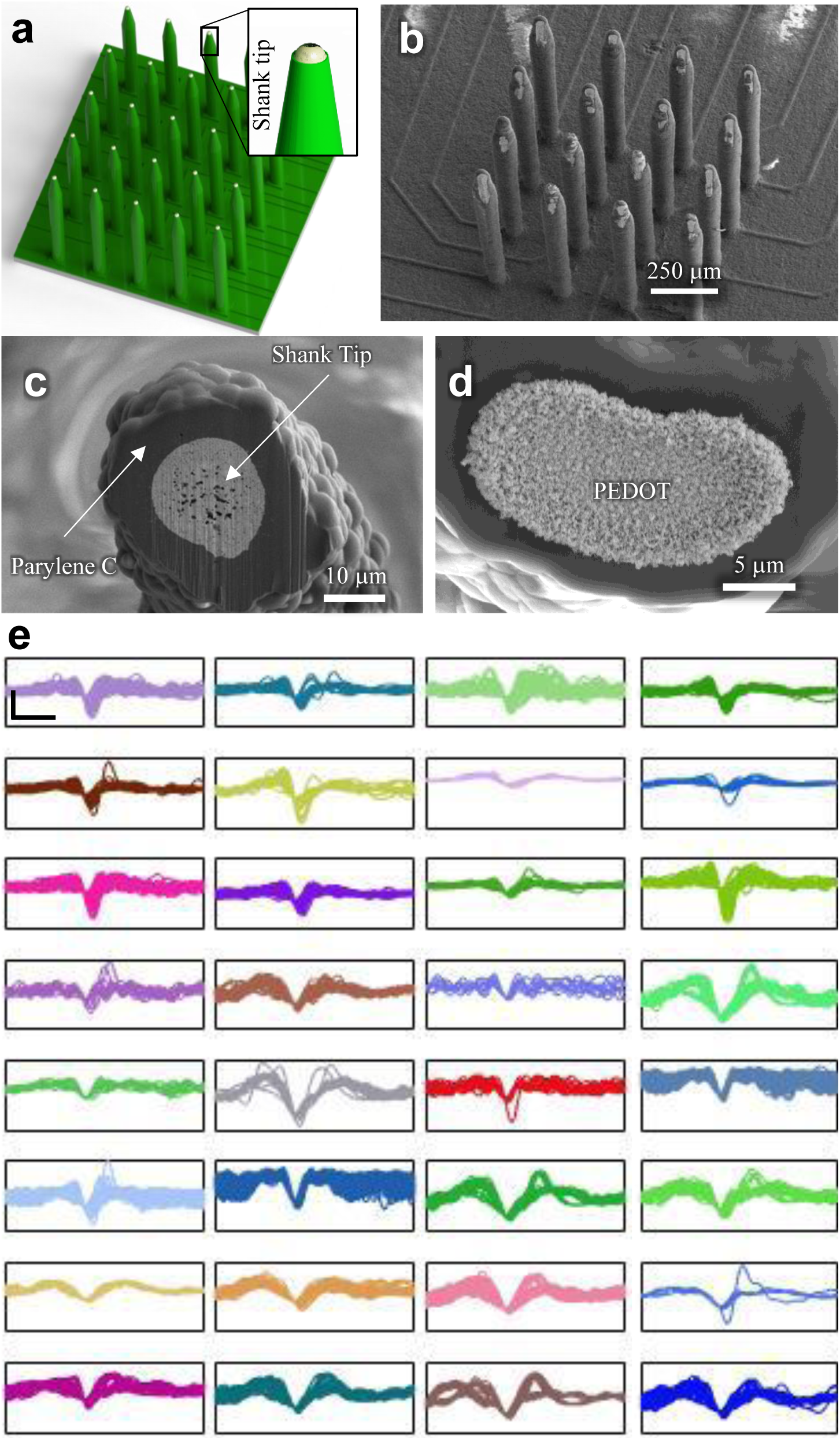
CMU Array functionalization and recording of action potentials from mouse brain. (a, b) A schematic and a SEM image of 3D printed probe with parylene insulation. (c) Exposure of the tip using focused ion beam, thus enabling the probe to record neural voltages. (d) Selective deposition of PEDOT at the tip using electroplating for reduced impedance. (e) Traces from 32 example channels of neural activity from one *in vivo* neural recording session in sensorimotor cortex of an anesthetized mouse. On some channels, waveforms from multiple neurons are identifiable. All data above a threshold of 1.5 SD are shown. Scale bar (top left) = 1 ms × 50 μV.

Probes were inserted into the anesthetized mouse brain (see Methods for recording details). We were able to record signals and isolate action potential waveforms from individual cortical neurons in sensorimotor cortex (depth = 660 μm). Each panel in Fig. 3e illustrates a different channel from a single session. These represent the first ever *in vivo* neural recordings using a 3D printed electrode. Signals from channels were recorded simultaneously sorted offline using a threshold set at four standard deviations beyond the median filtered signal (See Methods). On any given channel, we were able to identify individual action potential waveforms from 0-2 putative neurons (see Supplementary Fig. 3 for recording of two identifiable units on a single site). Mean firing rates of isolated neurons were comparable to that typically observed (mean 9.2 Hz, min/max 0.2-17.4 Hz with two putative multiunits at ∼40 Hz in the recording session shown). As an additional benchmark, we compared the quality of signal between the CMU Array and a top of the line commercial silicon MEA (Cambridge NeuroTech). Recordings from mouse motor cortex were performed in separate animals using the same data acquisition system and sorting analysis. The 3D printed array had an excellent signal to noise ratio (mean 7.5, SEM 0.6), and similar neuron yield per site as that found with commercial probes.

Although we are able to fabricate probes with high shank density, a creative, customizable back-end connectivity solution is necessary for it to be successful. In the next sections we show that the metal nanoparticle printing can be combined with polymer printing using the same aerosol jet 3D printer to develop a multi-layer custom PCB to route the signals from high-density shanks to an external amplifier. In addition, we also show that the probe can be directly printed on a commercially available pre-wired PCB. These results have strong implications spanning several device technologies in biomedical engineering.

### Multi-layer, Multi-material Custom Routing of Signals

As the high-density electrode leaves insufficient room for electrical leads, routing out the electric signals required an innovative solution. We again made use of the flexibility that 3D printing affords. We developed a multi-leveled, multi-material printing method to route the electrical signal to the appropriate recording devices (Fig. 4a). In the first step, a conductive metal layer (L1) of silver was printed on an alumina substrate as shown in Fig. 4b-i and the sintered in an oven. A layer of liquid polyimide polymer (L2) was then printed on the silver layer to form an insulating layer of 6 μm thickness as shown in Fig. 4b-ii. The layer L2 was printed such that the ends of the lines in L1 were exposed for future connections. The polymer printing was achieved by aerosolizing the liquid polyimide in the aerosol jet printer and printing the aerosol droplets onto the silver lines of L1. The polyimide was heated to facilitate polymerization, which formed the insulating layer L2. This process can be repeated as many times as is necessary (Fig. 4b-iii, 4b-iv). An additional polymer layer printed on the topmost metal layer (Fig. 4b-v) completed the multi-layer PCB to route the electric signals. The example in Fig. 4c shows routing from a 100-channel probe from a 2 mm × 2 mm area. The density and multi-level layout of the electrical connections from the shanks to the pads for the CMU Array in Fig. 4c are highlighted in a backscatter electron image in Supplementary Fig. 4.

**Figure 4.**
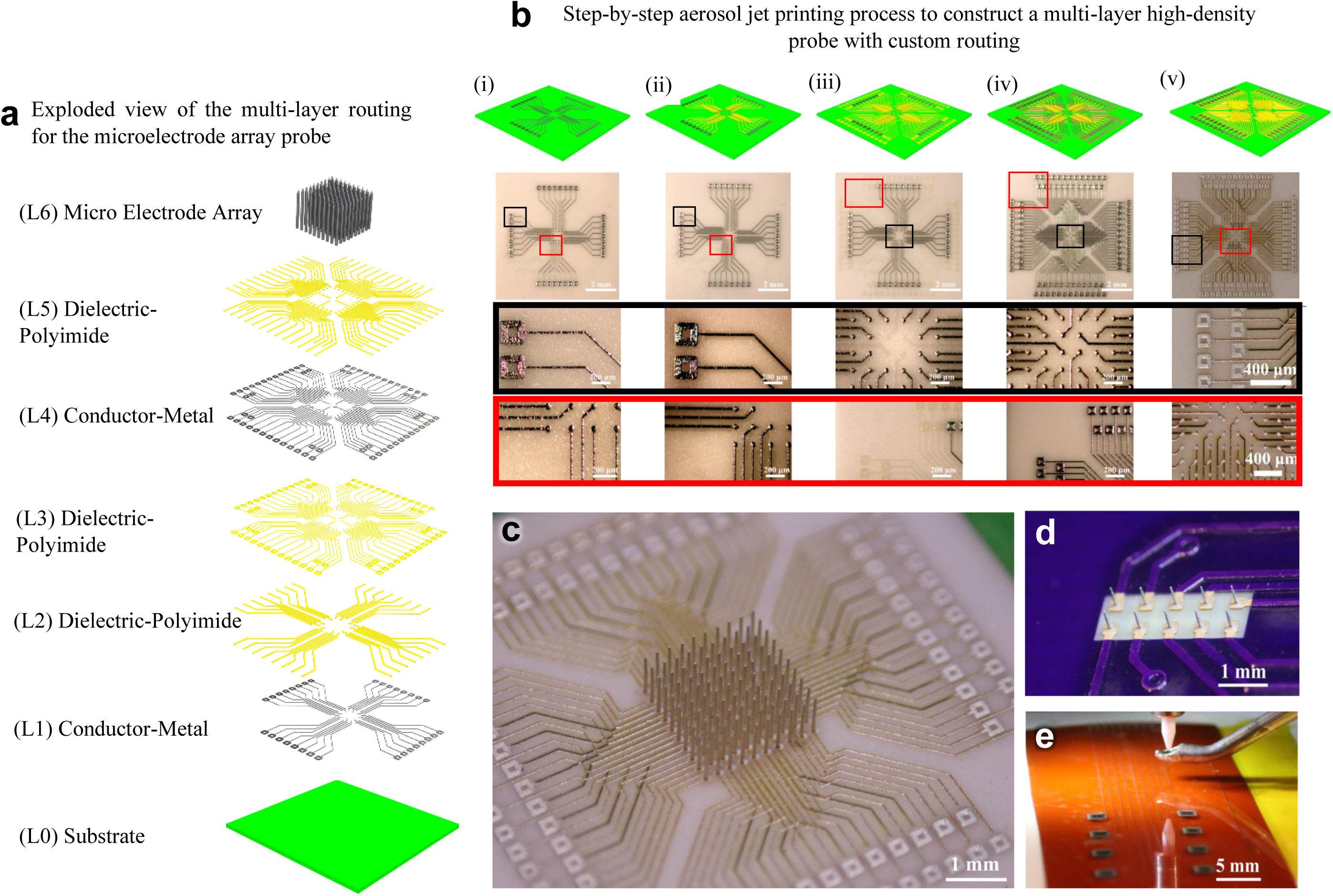
Electrical routing of the high-density shanks. (a, b) Multi-layer metal-polymer printing to fabricate a custom board to route signals from high density microelectrode array. (a) Exploded view of the different layers of the multi-layer probe. (b) Step-by-step process of printing the multi-layer board. (i) Printing of first metal layer on alumina ceramic with pads at both ends of each line. (ii) Polyimide printing to insulate the metal lines except at the pads. (iii) Middle polyimide layer to isolate the layer-4 from layer-1. (iv) Printing of the upper metal layer to create two conductive layers routing the signal out. (v) Polymer insulation printed on the upper metal layer except at the pads on the two ends. Note that the signal routing can be customized by simple changes to the printing program allowing arbitrary locations and heights for the probes and pads. (c) Image of a 100-shank probe over an area of 2 mm × 2 mm with printed multi-layer wiring/routing shown in (b). (d) 3D printed three dimensional microelectrodes on a commercially available PCB board which is pre-wired using lithographic processes. (e) Aerosol jet printed wiring on polyimide (Kapton) substrate.

In order to further exploit the flexibility offered by the nanoparticle printing process, we have also explored the printing of the shanks directly onto commercial PCBs, enabling additional flexibility in the construction of the bioelectrodes (Fig. 4d). The board had electroless nickel immersion gold (ENIG) metallization, a standard in the microelectronic industry. The micro-positioning accuracy for printing was within +/-1 μm. In order to enable probes on a flexible substrate shown in Fig. 1d, we also demonstrate the printing of wiring connections on a flexible polymer substrate (Kapton^®^ film, Fig. 4e). Our results establish that the aerosol jet printing offers enormous flexibility in routing the signals from the CMU Array demonstrated in this work.

## Discussion

The CMU Array demonstrated here overcomes current limitations in large-scale recording from a variety of biological tissue and opens up several new horizons for nanofabrication of biomedical solutions. First, the ability to print and route shanks at arbitrary locations allows the CMU Array to target specific regions of interest across distant areas of the brain – thus enabling precision science and reducing damage due to unwanted shanks inherent in one-size-fits-all probes. Indeed, increased study of the cascade of areas within the three-dimensional distribution of circuits in the brain appears to be the future of neuroscience.^15, 25-27^ The extremely high shank density we achieved (exceeding 6000 sites/cm^2^) represents an order of magnitude improvement over current microelectrode techniques. Second, as shown in Figs. 1 and 4, the metal-polymer printing developed in this paper allows rapid prototyping and custom routing of signals from shanks via changes to the printing programs. This combination of creating 3D objects (shanks) along with layered 2D planar wires (routing) will enable a rapid on-demand fabrication of study-specific probes or patient-specific neural interfaces. The use of flexible substrates provides an additional degree of customization not offered by the current methods. While extensive focus has been placed on increasing electrode channel counts, these channels are only helpful if they are able to be placed in the correct locations. The CMU Array constitutes a new avenue of targeted, optimized research that promises to reveal information processing strategies employed by neural ensembles across brain areas.^5^

The novel fabrication techniques also yield logistical improvements. Probe production time constitutes a matter of hours (followed by standard fab processes run in batch form the following day); rather than weeks. We note that aerosol jet printers with four nozzles that can print shanks simultaneously are commercially available and would further reduce the fabrication time. Lastly, we expect a significant reduction in the cost of the probes using this method, which will reduce the barriers to entry for researchers and clinicians that are either running hypothesis-driven experiments or aiming to create devices. Such ‘democratization’ of the microelectrode technology will immensely benefit researchers and clinicians.

We note that the proposed fabrication technique depends only upon aerosol droplet dynamics (Fig. 1a) and not on the nanoparticles inside the droplets. It is thus possible to construct the CMU Array from different materials, although the emphasis in this work largely has been on using silver due to the availability of the nanoparticle dispersions suitable for printing. The shanks can readily be plated with PEDOT (Fig. 4) and gold (Supplementary Fig. 5). We also note that although aerosol jet printing is used in the current work, other droplet based printing methods such as fine inkjet printing^28^ can be potentially explored construct the three dimensional shanks. Such an effort will be part of a future investigation.

The flexibility of production may provide further inroads across biomedical and material science. The printing of three-dimensional features provides a means to fabricate hollow metallic tubes. Such a construct can be used as microneedles for drug delivery and/or extraction of fluids; all while recording the bioelectric signals. The current method can also be combined with printing of three dimensional transparent polymers^19^ to construct optic-fiber paired probes for optogenetic stimulation and recording of neural signals. Lastly, the microelectrodes can be used in non-biological applications such as changing surface hydrophobicity through texturing, increasing energy storage in batteries through an increase in the surface area, and specific sensor devices.

In summary, we have demonstrated a rapid 3D additive printing method we developed to create a new class of customizable, high-density microelectrode array and demonstrated its superior operation in penetrating and recording from biological tissue. The technology increases recording sites per unit area by an order of magnitude and enables the on-demand, study-specific prototyping and manufacture of electrode configurations in a few hours. This technology paves the way to large-scale probes (thousands of channels; over several cm^2^ area) with easily modified probe layouts that can capture and potentially manipulate the dynamics of large, multi-area neural circuits with single-neuron and single-millisecond resolution.

## Acknowledgements

R.P.P. and E.A.Y. acknowledge support from the NIH (1R21EY029441) and the DSF Foundation. We thank the Andreas Pfenning group for providing the opportunity to test the probe in the macaque brain, and for Drs. Nick Melosh, Nick Anderson, Bin He, and Jack Beuth for helpful discussions as we prepared the study.

## Author contributions

M.S.S., E.A.Y., and R.P.P. conceived and designed the experiments. M.S.S., S.M.R., M.A.N., AND R.B. performed the experiments. J.W.R. and M.C. prepared the PEDOT plating recipe. All authors analyzed the data and discussed the results. M.S.S., E.A.Y., and R.P.P. co-wrote the paper.

## Methods

### 3D Nanoparticle Printing

To construct the nanoprinted array (CMU Array), we used Ag nanoparticle ink, i.e., Ag nanoparticles dispersed in a solvent (Prelect TPS 50 G2, Clariant Group, Frankfurt, Germany). Ag was chosen due to the commercial availability of the inks (i.e. dispersions) for this printing method.^1, 17^ The Ag nanoparticle size in the ink was 30-50 nm, the ink viscosity was about 1.5 cP, and the Ag particle loading in the ink was about 40 ± 2 wt %. The solvents used for this ink were de-ionized (DI) water and ethylene glycol. Ethylene glycol acts as a humectant and dispersant to help in the formation of the shank/pillar structures required for this work. Different substrates were used for this work. One of the substrates was alumina slab with 96% purity (ALN-101005S1, MTI Corp, Richmond, CA). The second substrate was silicon wafer. For alumina and Si substrates, the printing of silver nanoparticles and Kapton polymer were used to construct the leads to route the signal from the shanks. The third substrate was a PCB board with pads with dimensions of 340 × 340 μm and a metallization of electroless nickel immersion gold (ENIG). The microelectrodes were insulated using 3-5 microns of Parylene C (SCS Labcoter-2, dimer mass 8.3 g, furnace setpoint 690 °C, chamber gauge setpoint 135 °C, vaporizer setpoint 175 °C, vacuum setpoint 35 mTorr). Parylene C was chosen as the insulator material due to its superior biocompatibility, high resistivity, impermeability to biological species, and conformal deposition process. Deposited thin-film thickness was verified by including a calibration Si wafer alongside the 3D printed electrode array in the deposition chamber and measured using reflectometry (Nanometrics Nanospec 210XP). The metal at the tips of the shanks was exposed by removing Parylene C using a focused ion beam (FIB) machine (Model SMI3050SE, Seiko Instruments, Chiba Japan) that used a beam of Ga^+^ ions.

An Aerosol Jet 3D printer (Model AJ-300, Optomec, Inc., Albuquerque, NM) was used to print the microelectrode arrays. The AJ system at Carnegie Mellon University consists of three atomizers (two ultrasonic atomizers and a pneumatic atomizer), a programmable XY motion stage, a UV light source, and a deposition head. For the construction of the microelectrode arrays, we used one ultrasonic atomizer to aerosolize the silver ink. The aerosolized ink droplets were carried to a nozzle by a carrier gas (N_2_). Inside the nozzle, the droplets were focused toward the nozzle with the help of a sheath gas (also N_2_) to form a micro-jet. In case of printing of polymers we used pneumatic atomizer. The printing process was carried out by a continues flow of droplets which was diverted or resumed by movement of a shutter. The diameter of the nozzle was 150 μm, which is known to give rise to an aerosol stream having diameter of ∼10 μm.^17^ Before printing, the geometry of the conductive part was drawn in AutoCAD using a program in the software AutoLISP (AutoCAD 2015, Autodesk Inc., San Rafael, CA) and converted to a “prg” file compatible with the printer software. The shanks were printed by droplet-over-droplet in a method developed by the PIs, where semicircular printing was done to create a layer of about 20 μm thickness within a fraction of a second. The gas flow rate in the AJ machine during printing was about 25 sccm, while that for the sheath gas was about 50 sccm. Note that for initial prototyping, we have employed silver nanoparticles (rather than e.g. gold, for which, nanoparticle inks are readily available). We are aware of the cytotoxicity of silver but 1) the electrodes described here are for acute use, 2) all silver is concealed behind PEDOT:PSS, gold, and/or Parylene C, and 3) the ink used is yet another customizable feature of the array. Despite the potential for concern, vigorous, normal action potentials were able to be maintained throughout the duration of the recordings, lasting in the tens of minutes. No evidence of gross cell death was observed in the histological analyses.

### Sintering Conditions

The microelectrode array printed on alumina substrates were sintered at 250 °C for 6 hours at a ramp rate of 1 °C/min (for shanks <1.5mm long) and 0.3 °C/min (for shanks >1.5 mm long). For shanks on alumina, an alternate sintering temperature of 450 °C for 2 hours gave the same results. The microelectrode arrays printed on the Kapton substrate were sintered at 335 °C for 2 hours (same ramp rate as that for samples on alumina substrate). The microelectrode array printed on the PCB substrate were sintered at 195 °C for 2 hours (ramp rate of 1 °C/min). The sintering temperatures and times were chosen from PIs’ previous work on the sintering of silver nanoparticles.^17, 21^ The sintering was carried out in a programmable oven with controlled heating rates (Neytech Vulcan furnace, Model 3-550, Degussa-Ney Dental Inc., Bloomfield, CT). The printed Kapton for the multi-layer routing pattern was sintered at 330 °C.

### PEDOT:PSS Electrodeposition

PEDOT:PSS electrodeposition was carried out using a recipe optimized at CMU based upon a prior report.^29^ The electroplating solution was prepared by combining and mixing thoroughly 2 ml DI water and 41.24 mg polysodium styrene sulfonate (PSS) in a test tube. 2.2 μl of 3,4-ethylenedioxythiophene (EDOT) was added using a micropipette. This mixture was mixed thoroughly via shaking. The solution was sealed and refrigerated for 24 hours before use. Prior to electroplating, the solution was brought to room temperature and mixed again before electrodeposition. A Keithley 2401 sourcemeter as a current controlled power supply was used for the electrodeposition. All chemicals were purchased from Sigma Aldrich.

### Gold Electroplating

Gold electroplating was carried out using in Earthcoat Cyanide Free Gold Plating Solution (24K) heated to 60 °C on a hot plate with stirrer (Model MS300HS, Medline Scientific, Oxon, United Kingdom). The solution (filled to an adequate depth for submersion in a 1 L thick glass beaker) was agitated to mix well prior to probe immersion, but no agitation was used during plating to prevent damage to the device. The probe was cleaned before plating by submerging it in DI water (3 separate beakers used in sequence to reduce contamination). The probe was plated using Spa Plating (Bath, United Kingdom) MF Rectifier at a current of 0.1 A for 5 minutes, using a stainless-steel electrode. After plating, the rinsing process was repeated (using 3 beakers with fresh DI water) to remove any remaining plating solution. Energy dispersive X-ray spectroscopy (EDS) analysis was done to verify that the solution was plating gold without contamination.

### Compression Test for the Shanks

Shanks were printed onto an alumina substrate and tested under compression. The tests were carried out in a universal testing machine (Instron Corp, Newton MA) at a compression rate of 2mm/ min and equipped with a 20N load cell. A camera (Canon EOS Ti7 Rebel, Canon Corp, Tokyo, Japan) with a magnifying lens (model Tube TS-160, Infiniti Photo-Optical Company, Boulder CO) was used to record the compression behavior.

### Electrode insertion tests

Insertion testing was performed on the brains of adult C57/B6 mice. Anesthesia was administered via 2-3% inhaled isoflurane and was continued throughout the implantation. Craniomoties were performed to access the brain and the dura resected. Probes were dipped in Evans blue dye and then were then lowered through the craniotomy using a manual stereotax manipulator (RWD) at a rate of 100-200 μm/min. The probe was left in place for 5 minutes before being removed. Following several minutes following the removal of the probe, the mouse was euthanized with an overdose of isoflurane and decapitation. All procedures were approved by the Carnegie Mellon University Institutional Animal Care and Use Committee. Macaque insertion was performed in a similar manner on an adult animal recently euthanized for experimental purposes.

### Electrophysiology

In a different set of experiments, neural signals from were recorded from mouse sensorimotor cortex. The acute (i.e. not chronically implanted) recordings were performed in anesthetized mice (1.25-2% isoflurane) and lasted approximately 30 minutes. During recordings, the whiskers were manually manipulated to encourage activity. A reference wire shorted to ground resting on the acrylic head cap was also isotonically connected to the brain tissue. Recorded voltages containing neural signals were routed out through the custom routing board described to an Intan-based amplifier and data acquisition system (Whisper System, Neural Circuits LLC) running SpikeGLX (github.com/billkarsh/SpikeGLX). Spikes from each channel were sorted offline using a custom-built spike sorter that applied a high-pass filter to sort out spike frequency bands (300-5000 Hz). The amplitude discrimination threshold was set at four standard deviations above and below the mean of the recording segments. For each peak exceeding the threshold in magnitude, a three-millisecond putative waveform aligned on the absolute minimum of the waveform was then stored. The signal-to-noise ratio for the waveforms over a session was calculated as the peak-to-peak amplitude of the average waveform divided by the calculated voltage RMS. Values of greater than 1.5 were considered to be quality units that were easily discriminable from the underlying noise floor.

### Histology

For reconstruction of probe placement, the array was dipped in Evan’s Blue before implantation. Electrode tracks were imaged in the 40 μm slices, typically orthogonal to the axis of entry. Brain slices were placed on slides for imaging electrode tracks using a mounting medium that included a DAPI stain, which labels the nuclei of individual cells.

## Supplementary Information

**Supplementary Video 1**: A video showing a 4×4 CMU Array being built by aerosol jet printing. In this case, the electrodes are spaced 400 μm from each other and a have base diameter of 90μm. The diameter can be reduced gradually to 10-15 μm to create a taper via the AutoCAD program. The signal collection circuitry printed on the substrate allows perfect integration of the device in a single process. The captured video (at 45° angle) is slowed down by 10× for the purpose of clarity.

**Supplementary Video 2**: A time-lapse video of the construction of 1 512-electrode shank array with variable shank heights shown in Fig. 1g.

**Supplementary figure 1:**
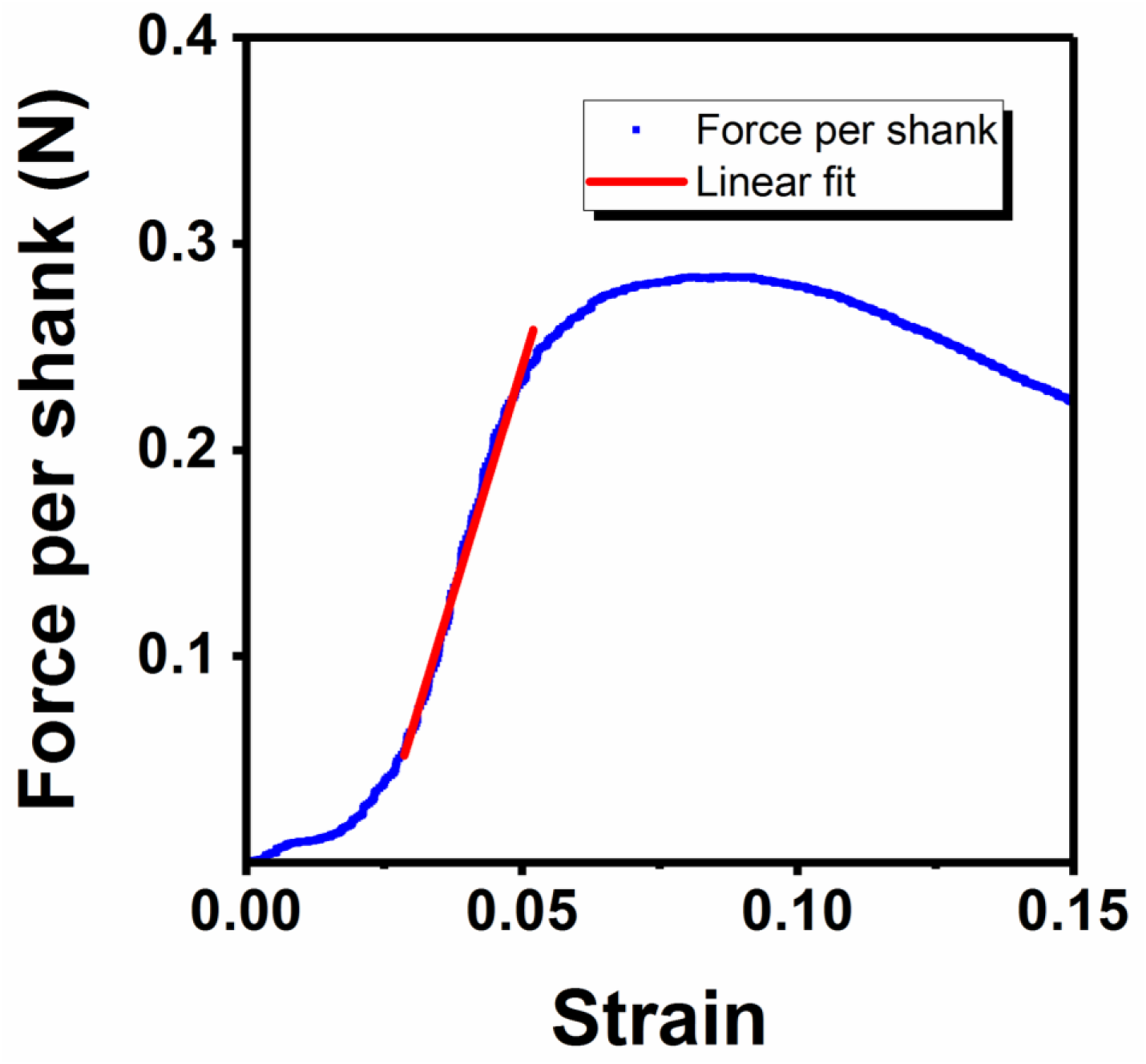
Load-displacement plot for the 3×3 array shanks with 90 μm diameter shown in Fig. 2a. The plastic deformation starts at about 7% strain. The shanks do not break until about 60% strain as shown in Fig. 2a. A larger diameter was chosen for this test due to the limitation in the compression testing apparatus.

**Supplementary figure 2:**
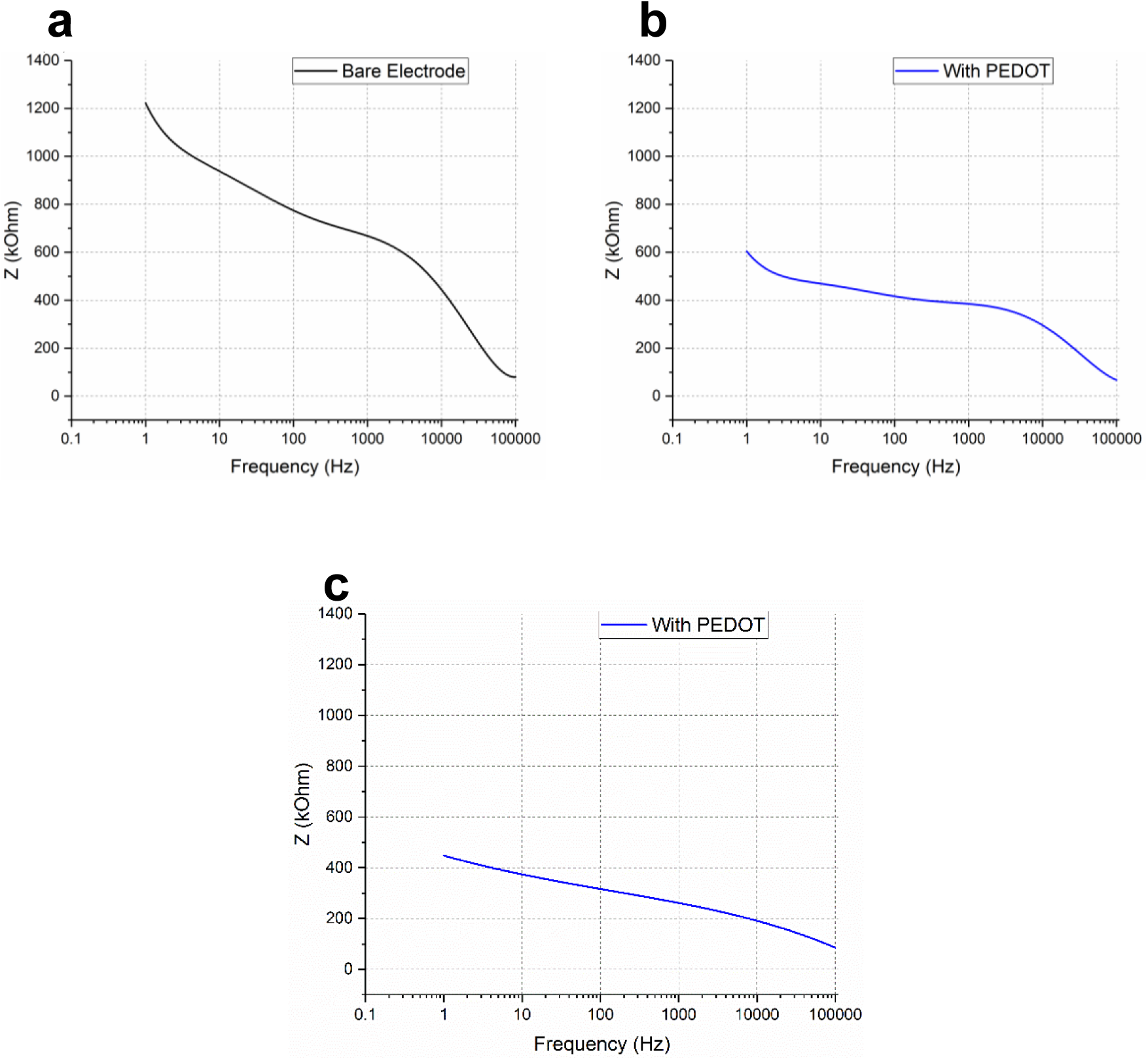
Electrical characterization of a CMU Array shanks showing an impedance in the range of 100s of kΩ at 1 kHz frequency. (a) An impedance of 662 kΩ at 1 kHz frequency without PEDOT coating. (b, c) Impedances of 379 kΩ and 256 kΩ at 1 Hz frequency for two different shanks with PEDOT coating.

**Supplementary figure 3:**
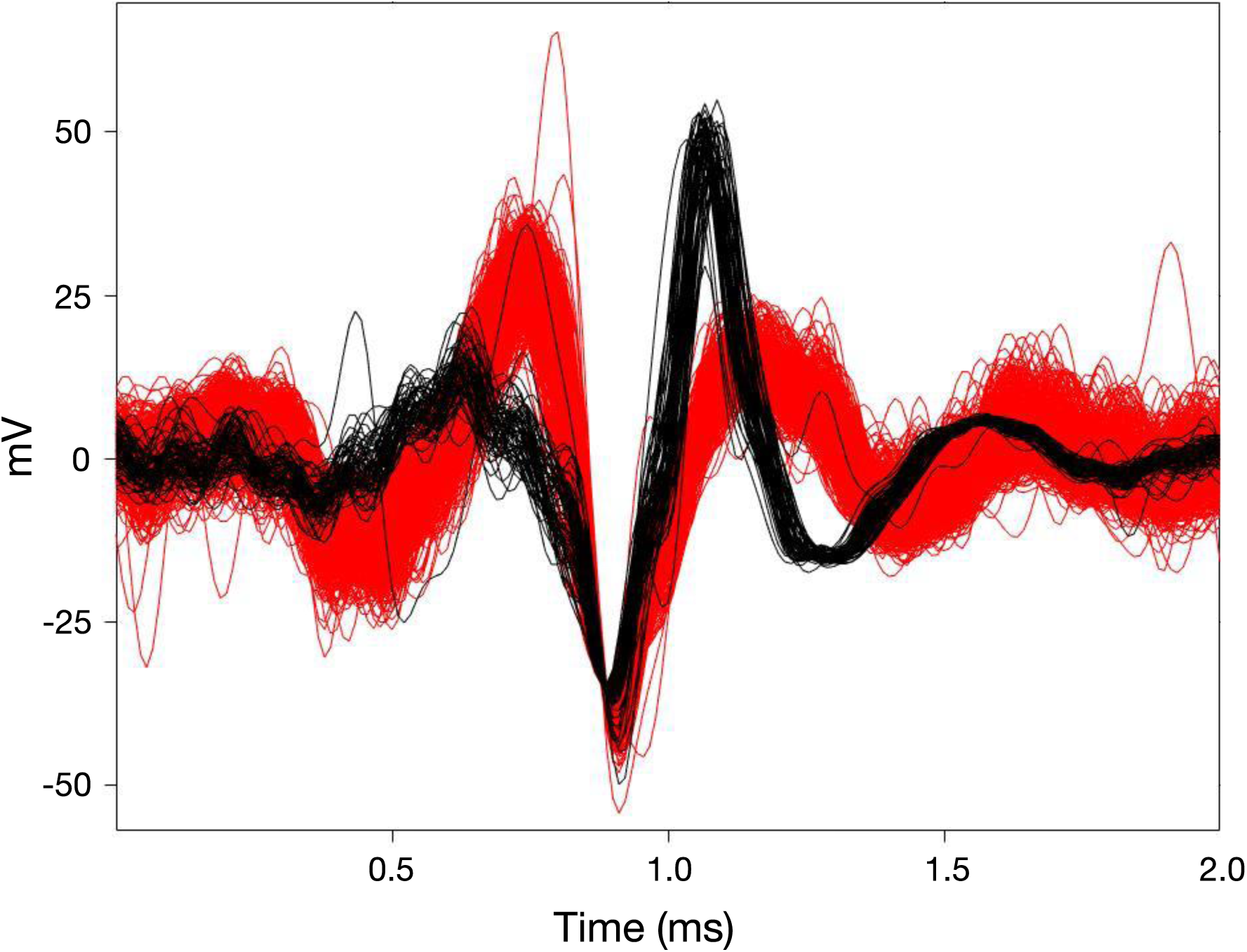
A representative channel with two easily distinguishable neuron waveforms, color-coded in black and red. Colors were applied using a basic envelope threshold to distinguish the between action potential waveforms of different amplitudes.

**Supplementary figure 4:**
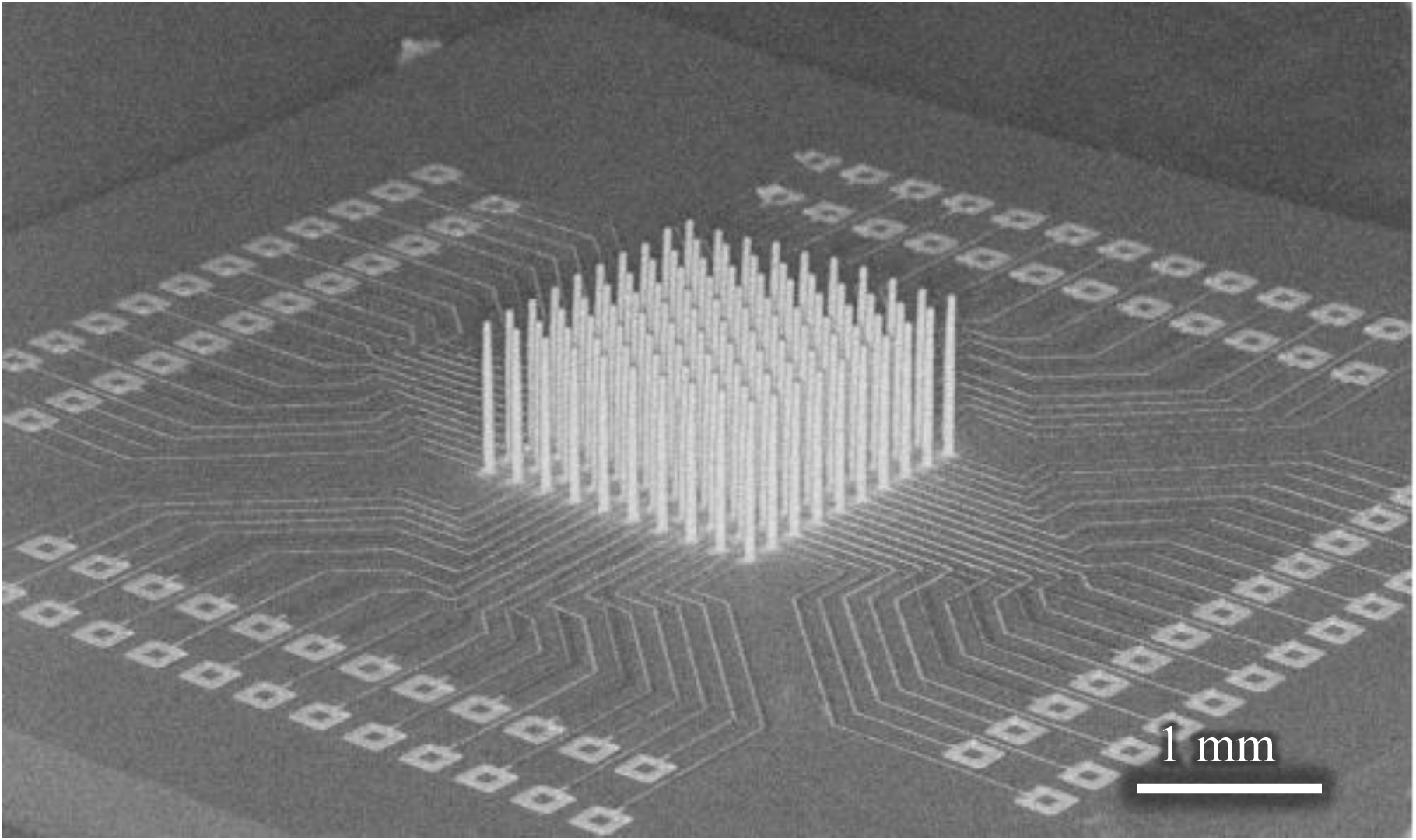
A SEM image (with backscatter electron mode) of the CMU Array shown in Fig. 4c. The routing complexity achieved by printing is evident.

**Supplementary figure 5:**
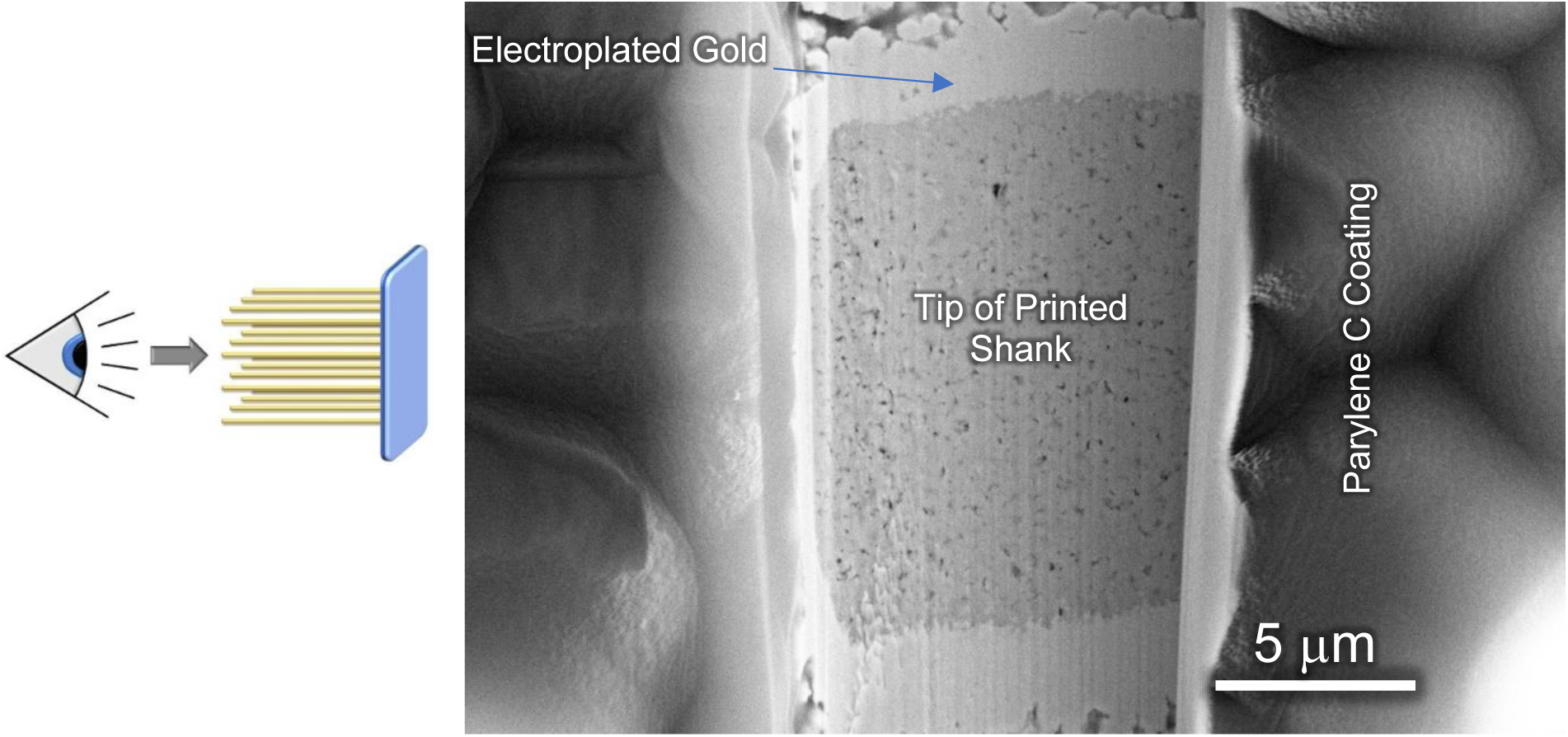
A Focused Ion Beam (FIB) section showing electroplated gold layer on 3D printed Ag shanks of the CMU Array (inset shows the orientation of the image). A standard cyanide-free gold plating recipe was used for the electroplating process (see Methods section of the paper).

